## supplementary information for "*Bacillus subtilis* RadA/Sms-mediated nascent lagging-strand unwinding at stalled or reversed forks is a two-step process: RadA/Sms assists RecA nucleation, and RecA loads RadA/Sms"

Running title: RecA and RadA/Sms work in concert

This supplementary file contains:

- Appendixes (A to G).
- Table (S1).
- Figures (S1 to S7).

**Appendix A**. *RadA/Sms does not form a stable complex with SsbA.* SsbA partially inhibits the maximal rate of ATP hydrolysis by RadA/Sms C13A. To test whether such inhibition is by a protein-protein interaction, we used a His-tagged RadA/Sms variant (1 µg) and native SsbA (1 µg). His-tagged RadA/Sms (predicted mass of 49 kDa) was retained in the 50-µl $Ni^{2+}$ matrix through coordination with its C-terminal histidine tag, and eluted (E) with buffer A containing 400 mM imidazole (Figure S2A, lanes 3-6). However, SsbA (predicted mass 18.6 kDa) was mainly present in the flow-through (FT), and the traces of the protein entrapped in the $Ni^{2+}$ matrix were released after the first wash (W1) with buffer A (Figure S2A, lanes 7-10). When both, His-RadA/Sms and SsbA, were loaded into the $Ni^{2+}$ column, we observed that SsbA was present in the FT (Figure S2B, lane 3), while His-RadA/Sms was observed in the E fraction in the presence of 400 mM imidazole (Figure S2B, lane 6). The same result was observed when His-RadA/Sms and SsbA were incubated in the presence of 10 µM cssDNA and 5 mM ATP (Figure S2B, lanes 7-10). This suggests that RadA/Sms does not form a stable complex with SsbA.

**Appendix B**. *RadA/Sms K104R or RadA/Sms K255R does not affect the PcrA ATPase.* RadA/Sms K104R or RadA/Sms K255R interacts with and blocks the ATPase of RecA (Figure 1C-D). Similarly, *wt* RadA/Sms interacts with and inhibits the ATPase of RecA [1]. To test if this inhibition is specific, the effect of RadA/Sms over the ATPase activity of an unrelated enzyme was measured. PcrA is a highly efficient ssDNA-dependent ATPase that physically interacts with RecA [2]. RadA/Sms K104R or RadA/Sms K255R (1 RadA/Sms variant/330-nt) was incubated with limiting concentrations of PcrA in buffer A containing 5 mM ATP and an ATP regeneration system [3]. The ssDNA-dependent ATPase of PcrA ($V_{max}$ 25.8 ± 4.51 µM · $min^{-1}$) was not affected by the presence of RadA/Sms K104R or RadA/Sms K255R when compared with PcrA alone (Figure S3B). Similar results were observed when *wt* RadA/Sms was used as a control. Thus, we can rule out any non-specific effect of RadA/Sms K104R or RadA/Sms K255R on the ATP hydrolysis rate of RecA.

**Appendix C**. *RadA/Sms and its mutant variants provide a roadblock rather than reversing RecA-mediated DNA strand exchange.* We have tested whether RadA/Sms and its mutant variants RadA/Sms C13A (unable to interact with RecA) and RadA/Sms K104A (unable to bind ATP) affect homology-directed RecA-mediated DSE using a three-strand recombination assay. In the mock reaction, RecA, in concert with the two-component mediator (SsbA-RecO), efficiently catalysed formation of joint molecule (*jm*) intermediates between the 3,199-nt cssDNA ([*css*], 10 µM in nt) and the complementary strand of the 3,199-bp dsDNA ([*lds*], 10 µM in bp) substrate. This is followed by extension of the heteroduplex region leading to nicked circular (*nc*) duplex and linear ssDNA products (Supplementary Figure S4A, lane 2). This reaction was catalysed even in the absence of an ATP-regeneration system [4].

Hexameric RadA/Sms·ATP or RadA/Sms C13A·ATP binds ssDNA and dsDNA with an apparent dissociation constant of ~2 nM and ~5 nM, respectively, but RadA/Sms K104A does it with about two-fold higher affinity [1,5]. In the presence of limiting to stoichiometric RadA/Sms or RadA/Sms C13A concentrations (0.8 to ~3 RadA/Sms/cssDNA molecule), the yield of RecA-mediated DSE (or *jm* + *nc* product formation) was not significantly affected ($p$ >0.1) in a 60 min reaction at 37 °C (Supplementary Figure S4A, lanes 3-5 and 8-10). The presence of ~3 RadA/Sms K104A /cssDNA molecule (at a RadA/Sms K104A:RecA 1:120 molar ratio) marginally reduced RecA-mediated DSE (Supplementary Figure S4A, lane 15). Addition of ~6 RadA/Sms, RadA/Sms C13A or RadA/Sms K104A /cssDNA molecule (at a 1:60 stoichiometry), significantly reduced ($p$ <0.01) RecA-mediated DSE (Figure S4A, lanes 6-7, 11-12 and 16-17). It is likely that: i) few RadA/Sms [or its mutant variants]/cssDNA or dsDNA molecule limit homology-directed RecA-mediated DSE on plasmid-sized DNA substrates; ii) such effect is independent of an active RecA disassembly from the cssDNA *via* a protein-protein interaction, because RadA/Sms C13A, which fails to interact with RecA, limits DSE; and iii) this inhibition is not due to a displacement of the invading strand by RadA/Sms, because RadA/Sms K104A, which cannot unwind DNA, inhibits RecA-mediated DSE. *E. coli* mutant variants equivalent to RadA/Sms C13A and RadA/Sms K104A ($RadA_{Eco}$ C28Y or $RadA_{Eco}$ K108R) also reduce the efficiency of DSE [6]. In contrast, *wt* $RadA_{Eco}$, which seems to lack DNA helicase activity, acts as a *mediator* and accelerates the initial step of DSE to yield the final $RecA_{Eco}$-mediated products more quickly without changing the final efficiency of product formation, at a 1:17 stoichiometry relative to RecA and in the absence of $RecO_{Eco}$ [6].

We can envision that RecO, by promoting SsbA displacement from the displaced strand and aiding end-dependent RecA nucleoprotein filament disassembly, indirectly compromises the DSE reaction. To test the hypothesis, the three-strand exchange reaction was performed in the presence of dATP, since Firmicutes RecA·dATP catalyses DSE in the absence of the two-component mediator (SsbA and RecO) *in vitro* [4,7-9]. However, since $RadA_{Eco}$ does not facilitate RecA-mediated DSE when $SSB_{Eco}$ was omitted from the reaction [6], and for the sake of comparison, SsbA was added. Here, RecA-mediated DSE was significantly reduced and inhibited ($p$ <0.01) in the presence of ~6 and ~12 RadA/Sms, RadA/Sms C13A or RadA/Sms K104A /cssDNA molecule (Supplementary Figure S4B, lanes 6-7, 11-12 and 16.17 *vs*. 2). It is likely that the three hexameric RadA/Sms proteins limit RecA-driven DSE acting as a roadblock upon binding to cssDNA and/or to a recombination intermediate *in vitro*. Alternatively, RadA/Sms may reverse RecA-mediated DSE.

To discriminate whether RadA/Sms reduces the efficiency of RecA-mediated DSE, or reverses DSE, the *css* and *lds* DNA substrates were incubated with fixed concentrations of SsbA, RecO and RecA for a variable time (10, 20, 30, 45 or 60 min) to allow the formation of intermediates. Then, ~6 RadA/Sms /cssDNA molecule were added and the reaction incubated further (60 min at 37 °C). Independently of the time point at which RadA/Sms is added, the reaction does not progress further when compared to the absence of RadA/Sms (Figure S4C, lanes 4, 7, 10, 13 and 16 *vs* 3, 6, 9, 12 and 15). The amount of recombination intermediates and products is that accumulated before RadA/Sms addition (Supplementary Figure S4C, lanes 4, 7, 10, 13 and 16 *vs* 2, 5, 8, 11 and 14). Thus, RecA-mediated strand transfer reaction is limited, but the reaction is not driven backwards to the original substrates.

**Appendix D**. *Defining the length of RadA/Sms binding site.* The hexameric RadA/Sms DNA helicase efficiently unwinds a 5′-tailed duplex substrate with a 5′→3′ polarity. Previous assays were performed with substrates with ssDNA tails ≥30-nt [1,10]. To test for the minimal length required for efficient RadA/Sms-mediated DNA unwinding, tailed duplex substrates with a variable length of the 5′-tail (from 5- to 30-nt) were used (Figure S1). As revealed in Supplementary Figure S5, RadA/Sms (20 nM) did not unwind a blunted duplex or 5′-tailed duplexes up to 10-nt long tails. A 15-nt tailed substrate was very poorly unwound, and the maximal rate of RadA/Sms-mediated unwinding was observed using a duplex substrate with a 30-nt long tail.

**Appendix E**. *RecA loads RadA/Sms to unwind a HJ-Lead DNA substrate.* Different scenarios (*a* to *g*) can be envisioned to explain how RecA activates RadA/Sms unwinding of the nascent lagging-strand of a HJ-Lead DNA substrate. In condition (*a*), traces of an exonuclease contaminating the RecA protein preparation would degrade the 3′-end of the nascent leading-strand to free a 5′-tail and facilitate RadA/Sms assembly on the nascent lagging-strand. This hypothesis is unlikely because: i) the nascent leading-strand is the radiolabelled strand, and its degradation was not observed (Figure 3A, lane 1 *vs*. 6-8); and ii) RadA/Sms C13A should also unwind the HJ-Lead substrate, but this was not observed (Figure 3C).

In condition (*b*), dynamic RecA association/dissociation may partially open the duplex DNA at the junction to generate a RadA/Sms entry site. To test the hypothesis, a 5′-fork DNA with target sites for overlapping restriction enzymes, at the duplex region, was constructed (Supplementary Figure S7A). As revealed in Appendix F, RecA bound to the poorly hydrolysable ATP analogue (ATPγS) RecA·ATPγS polymerised on the 3′-tail of 5′-fork DNA may partially open the duplex DNA at the junction, as judged by its sensitivity to the KpnI enzyme, and this could generate a RadA/Sms entry site, but RecA·ATP does not (Supplementary Figure S7B). Furthermore, RecA cannot facilitate the recruitment of RadA/Sms C13A onto the nascent lagging-strand, confirming that a direct protein-protein interaction is necessary and sufficient to activate RadA/Sms unwinding of the nascent lagging-strand.

In condition (*c*), hexameric RadA/Sms may interact with and activate RecA to remodel a HJ-Lead substrate, as proposed for RecA$_{Eco}$ [11]. We consider this option unlikely RecA does not remodel a stalled or reversed fork (Figures 3A, lane 3 and 5A-B, lane 3). Moreover, RecA$_{Eco}$-mediated fork reversal exhibits an absolute requirement for ATP hydrolysis [11]. However, the RadA/Sms variants RadA/Sms K104A or K225R interacts with and blocks RecA-mediated ATP hydrolysis (Figure 1A, C-D), and RadA/Sms unwinds the nascent lagging-strand in the presence of preformed RecA·ATPγS nucleoprotein filaments (Appendix F, Supplementary Figure S7C, lane 7). In addition, RadA/Sms cannot activate RecA to remodel a fully replicated fork (3′- 5′-fork DNA, a replicating fork with fully synthesised leading- and lagging-strands, Figure S1) (Appendix G, Supplementary Figure S7D).

In condition (*d*), RecA bound to the 3′-tailed leading-strand frays the nascent lagging-strand, and this is sufficient to allow RadA/Sms self-recruitment to the end of the nascent lagging-strand to unwind the HJ-Lead DNA substrate. We did not favour this option because RadA/Sms C13A is not able to bind to the hypothetical access provided by RecA (Figure 3C), further confirming that a direct RecA-RadA/Sms interaction is necessary.

In condition (*e*), RecA interacts with RadA/Sms monomers in solution to construct and deposit a hexamer around the nascent lagging-strand of the HJ-Lead DNA to unwind the DNA substrate. We consider this hypothesis unlikely, because RadA/Sms and its variants (RadA/Sms C13A, RadA/Sms K104A or RadA/Sms K255R) are hexamers in solution [1,10], and as postulated by the ring-maker strategy [12] discrete RadA/Sms subunits would be needed.

In condition (*f*), RadA/Sms, which belongs to the LonA superfamily, may adopt an hexameric open-spiral structure configuration, as reported for the LonA$_{Eco}$ protease [13]. RecA bound to the 3′-tailed HJ-Lead DNA may interact with, load and close the RadA/Sms ring to unwind the nascent-lagging strand by the ring-closure strategy. We consider this option unlikely because RadA/Sms forms a closed hexameric ring in solution with two-fold symmetry and three protomers per symmetric unit [10,14] rather than a right-handed open spiral.

In condition (*g*), monomeric or hexameric RecA, at a 6:1 or 1:1 stoichiometry, respectively, relative to RadA/Sms, upon binding to and extending the ssDNA [15] may physically open a hexameric RadA/Sms ring and deposit it onto the nascent-lagging strand of the HJ-Lead DNA with RadA/Sms unwinding the substrate as proposed by the ring-breaker strategy [16]. We favour this strategy, although the precise molecular mechanism used by RecA to open the ring-shaped hexamer and activate RadA/Sms to unwind a HJ DNA substrate remains unknown.

**Appendix F**. *RecA-mediated ATP hydrolysis is not required for RadA/Sms recruitment at the nascent lagging-strand.* To explore condition (*b*), a 5′-fork DNA substrate whose nucleotide sequence contains three restriction sites overlapping by 2-bp at the parental 30-bp long duplex segment was constructed (Supplementary Figure S7A). This 5′-fork DNA was pre-incubated with increasing RecA (100-400 nM) concentrations in the presence of ATP or ATPγS as cofactors (15 min at 37 °C). Then, the reaction mixture was incubated with the KpnI, SmaI or BamHI restriction enzymes (60 min at 37 °C), and the products visualised by PAGE.

The polarity of RecA filament disassembly from ssDNA, which occurs predominantly from filament ends, relies on ATP hydrolysis [17,18]. RecA·ATP, nucleated on the 5′-fork DNA away from the junction in the 5′→3′ direction, did not promote any significant opening of the duplex, as judged by the sensitivity of the 3′-fork DNA to the three (KpnI, SmaI and BamHI) restriction enzymes (Supplementary Figure S7B). This suggests that its secondary DNA-binding site bound to the complementary strand may not open it. Unexpectedly, when RecA·ATPγS was tested, ~90%, ~50% and ~35% of the 5′-fork DNA substrate was resistant to KpnI, SmaI and BamHI cleavage, respectively, in the presence of 400 nM RecA·ATPγS (Supplementary Figure S7B). Thus, it is likely that a RecA·ATPγS filament polymerizing toward the branch point partially opens the duplex DNA, as judged by its resistance to restriction enzyme cleavage.

In the presence of 400 nM RecA·ATPγS, only ~50% of the substrate is resistant to BamHI digestion (Supplementary Figure S7B), translating in duplex opening of up most to 17-nt (Supplementary Figure S7A). Since RadA/Sms requires a longer ssDNA region for self-loading (Supplementary Figure S5), we assumed that the interaction with RecA should be strictly necessary for RadA/Sms activation. Then, we have tested whether fixed RadA/Sms can unwind the non-cognate HJ-Lead DNA substrate in the presence of increasing RecA·ATPγS concentrations in buffer A containing 2 mM ATP (15 min at 37 °C). As revealed in Supplementary Figure S7C, a RecA·ATPγS filament was unable to separate the paired strands (lane 3). A preformed RecA·ATPγS nucleofilament interacts with and loads RadA/Sms on the nascent lagging-strand to unwind the substrate with similar efficiency than in the presence of RecA·ATP (Figure 2A, lanes 6-8 *vs*. Supplementary Figure S7C, lanes 5-7), suggesting that a RecA·ATPγS nucleofilament interacts with and recruits RadA/Sms to the nascent lagging-strand.

**Appendix G**. *RadA/Sms cannot activate RecA to remodel a fully replicated fork.* To test whether RadA/Sms interacts with and activates RecA to remodel a fully replicated fork, an artificial substrate (3′-5′-fork DNA) was constructed (Supplementary Figure S1). To confirm the integrity of this DNA substrate, the PcrA enzyme was used. As previously shown [2], PcrA binds and translocates on the template lagging-strand to generate a 5′-tailed intermediate and the radiolabelled strand (Supplementary Figure S7D, lane 13).

In the presence of a fixed RecA (400 nM) concentration the remodeling of 3′-5′-fork DNA was not observed (Supplementary Figure S7D, lane 5). Similarly, increasing concentrations of RadA/Sms (10 – 80 nM) or fixed RadA/Sms (20 nM) and increasing RecA (100 – 400 nM) concentrations did not remodel the substrate (Supplementary Figure S7D, lanes 6-12). It is likely that: i) a large excess of RecA (800 RecA monomers/3′-5′-fork DNA molecule) fails to remodel a fully replicated fork; and ii) RadA/Sms fails to activate RecA to remodel a 3′- 5′-fork DNA substrate.

**Table S1**. RecA- and RadA/Sms-mediated ATP hydrolysis

| DNA substrate and additional protein | $V_{max}$ (μM · min$^{-1}$) | | | | |
|---|---|---|---|---|---|
| | RecA | RadA/Sms | RadA C13A | RadA K104R | RadA K255R |
| none | <0.1 | 1.9 ± 0.1 | 1.9 ± 0.1 | <0.1 | <0.1 |
| cssDNA[a] | 7.68 ± 0.3 | 1.9 ± 0.2 | 9.8 ± 0.3 | <0.1 | <0.1 |
| cssDNA + SsbA | 0.4 ± 0.1 | 1.9 ± 0.1 | 4 ± 0.2 | - | - |
| cssDNA + RecO + SsbA | 14.4 ± 0.2 | - | - | - | - |

| DNA substrate and additional protein | $V_{max}$ (μM · min$^{-1}$) | | | |
|---|---|---|---|---|
| | RadA + RecA | RadA C13A + RecA | RadA K104R + RecA | RadA K255R + RecA |
| cssDNA | 4.0 ± 0.3 | 20.3 ± 0.2 | 1.1 ± 0.1 | 1.0 ± 0.1 |
| cssDNA + SsbA | 4.2 ± 0.3 | 6.3 ± 0.3 | <0.5 | <0.5 |
| cssDNA + SsbA + RecO → RecA + RadA | - | | 1.2 ± 0.2 | 1.1 ± 0.2 |
| cssDNA + SsbA + RecO + RecA→ RadA | - | | 5.2 ± 0.3 | 4.4 ± 0.3 |
| cssDNA + RecA + RadA → SsbA + RecO | - | | 1.1 ± 0.1 | 1.1 ± 0.1 |

The 3,199-nt cssDNA (10 μM in nt) was incubated with the indicated protein(s), and the rate of ATP hydrolysis was measured as described (see Materials and methods). When indicated, *wt* RadA/Sms was replaced by RadA/Sms C13A, K104R or K255R variants. In some conditions, SsbA (150 nM) and RecO (100 nM) were added. The steady-state kinetic parameters for RadA/Sms (30 nM), RecA (800 nM) or both proteins were derived from more than three independent experiments like those shown in Fig. 1A and 1C-D. The results are shown as mean ± SEM. The + symbol indicates the proteins added, and arrows denote that the ssDNA was incubated (5 min at 37 ºC) with the preceding protein(s) to preform cssDNA-protein complex(es). ATP hydrolysis was then monitored. -, not done or it does not apply.

Figure S1

| DNA | Substrate description | Structure + oligonucleotide composition |
|---|---|---|
| HJ-Lead | 3'-tailed HJ DNA | 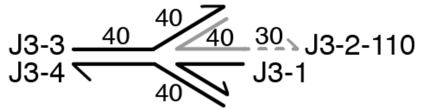 |
| HJ-Lag | 5'-tailed HJ DNA | 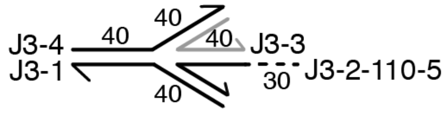 |
| forked-Lead | leading-strand-gapped forked DNA | 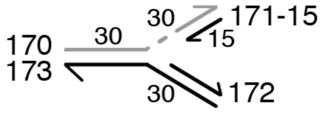 |
| forked-Lag | lagging-strand-gapped forked DNA | 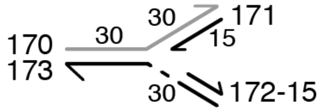 |
| 5 duplex | 5-nt 5'-tail duplex DNA | 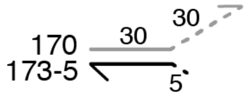 |
| 10 duplex | 10-nt 5'-tail duplex DNA | 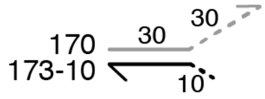 |
| 15 duplex | 15-nt 5'-tail duplex DNA | 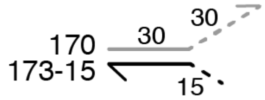 |
| 20 duplex | 20-nt 5'-tail duplex DNA | 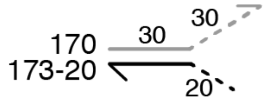 |
| 30 duplex | 30-nt 5'-tail duplex DNA | 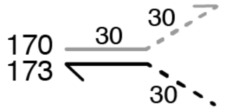 |
| 5'-fork-RE | 5'-fork DNA with restriction enzyme sites | 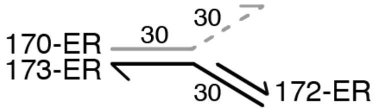 |
| fork | fully replicated fork | 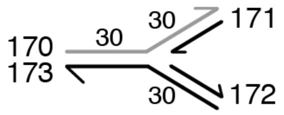 |

Figure S1. DNA substrates used in the assays. The DNA substrates were constructed by annealing the indicated oligonucleotides. In grey is denoted the radiolabelled strand, and in dotted lines the noncomplementary regions.

Figure S2

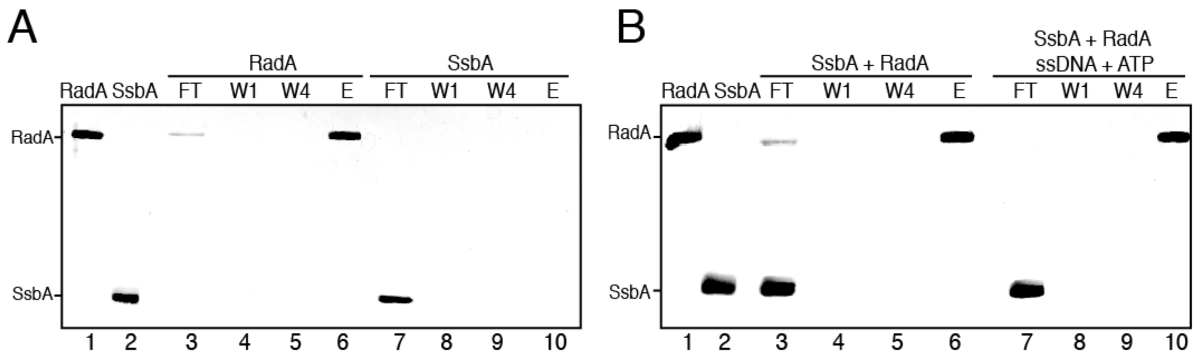


Figure S2. RadA/Sms does not form a stable complex with SsbA. His-tagged RadA/Sms (1 µg) and/or native SsbA (1 µg) was (were) incubated or not with 10 µM cssDNA and 5 mM ATP (5 min, 37 ºC) in buffer A. The pre-incubated mixture was loaded onto a 50 µl $Ni^{2+}$ matrix (5 min, 37 ºC), and the flow-through (FT) was collected. The $Ni^{2+}$ matrix was washed (W) four times with four column volumes of buffer A containing 20 mM imidazole, and the retained proteins were finally eluted (E) with buffer A containing 0.4 M imidazole. The proteins were separated by 10% SDS-PAGE, and gels were stained with Coomassie Blue. In lanes 1 and 2, the proteins (1 µg) were loaded for evaluating retention on the column. The experiment was repeated three times with similar results, and a representative gel is shown here.

Figure S3

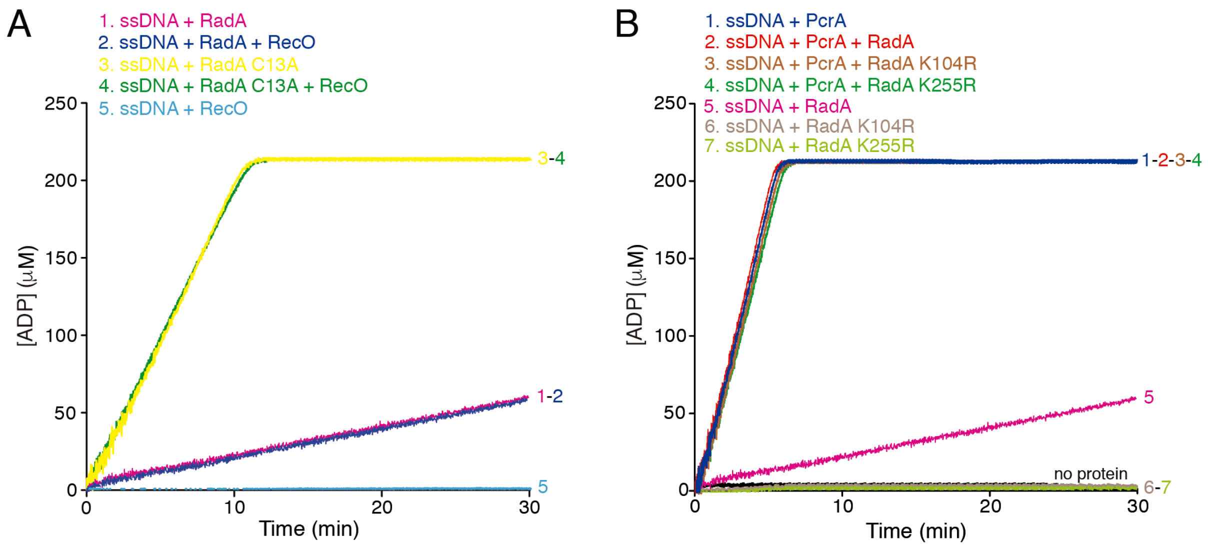


Figure S3. RecO does not affect RadA/Sms-mediated ATP hydrolysis (A) and RadA/Sms does not affect PcrA-mediated ATP hydrolysis (B). The cssDNA (10 µM in nt) was incubated with the indicated protein(s) (*wt* RadA/Sms or its C13A, K104R or K255R variants [30 nM], PcrA [15 nM], and RecO [100 nM]) in buffer A containing 5 mM ATP and the ATP regeneration system. The ATPase activity was measured for 30 min. All reactions were repeated three or more times with similar results, and a representative graph is shown here.

Figure S4

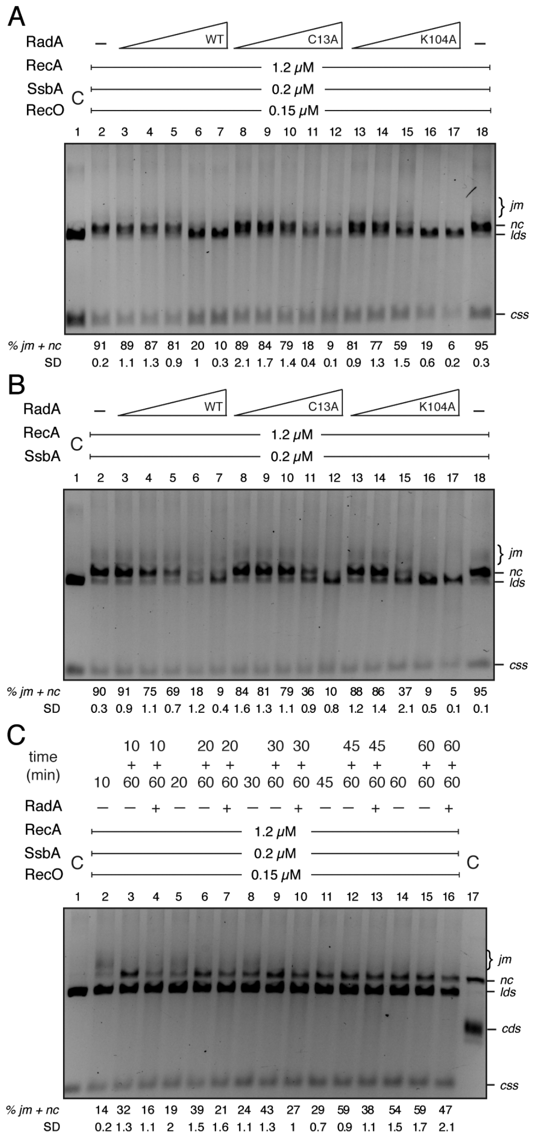


**Figure S4**. RecA-mediated DSE in the presence of *wt* and RadA/Sms mutant variants. (A) The 3,199-nt ssDNA (*css*) (10 μM in nt) and 3,199-bp KpnI-linearised dsDNA (*lds*) (10 μM in bp) were incubated with SsbA, RecO, RecA and *wt* RadA/Sms, RadA/Sms C13A or RadA/Sms K104A (2.5 to 40 nM) in buffer A containing 5 mM ATP (60 min at 37 ºC), and products were separated by 0.8% AGE. (B) The *css*and *lds* DNA substrates were incubated with SsbA, RecA and increasing *wt* RadA/Sms (or RadA/Sms C13A or RadA/Sms K104A) (2.5 to 40 nM) for 60 min at 37 ºC in buffer A containing 5 mM dATP, and products were separated by 0.8% AGE. (C) The *css* and *lds* DNA substrates were pre-incubated with SsbA, RecO, RecA for a variable time (10 - 60 min) in buffer A containing 5 mM ATP. Then, RadA/Sms (20 nM) was added or not, the reaction further incubated (60 min at 37 ºC), and products were separated by 0.8% AGE. In lane 1, the *css* and *lds* substrate were run as a control. The symbols - denote the absence of the indicated protein(s). The position of the bands corresponding to *css*, *lds*, *jm* and the *nc* product is indicated. Amounts of recombination intermediates plus product are indicated, and expressed as percentage of total substrate added. The results are the average of more than three independent experiments ± SD.

Figure S5

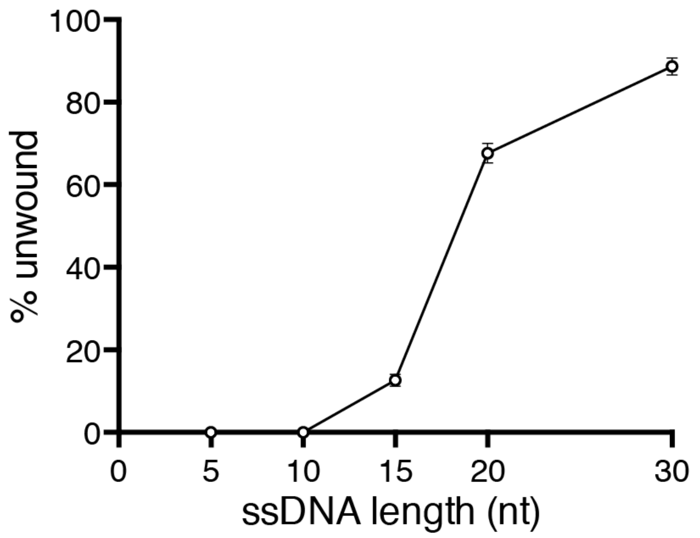


Figure S5. Relationship between the 5′-tail length of DNA and RadA/Sms-mediated DNA unwinding. [γ-$^{32}$P] 5′-tailed duplex DNA substrates (0.5 nM) with increasing tail lengths (5-10-15-20-30 nt) were incubated with RadA/Sms (20 nM) in buffer A containing 2 mM ATP for 15 min at 30 ºC. Reactions were stopped, deproteinised and separated by 6% native PAGE. Gels were dried and visualised by phosphor imaging. The fraction of unwound products in three independent experiments was quantified, and the mean ± SD is represented here. See Figure S1 for details of substrate composition (substrates denoted as 5 duplex, 10 duplex, 15 duplex, 20 duplex, 30 duplex).

Figure S6

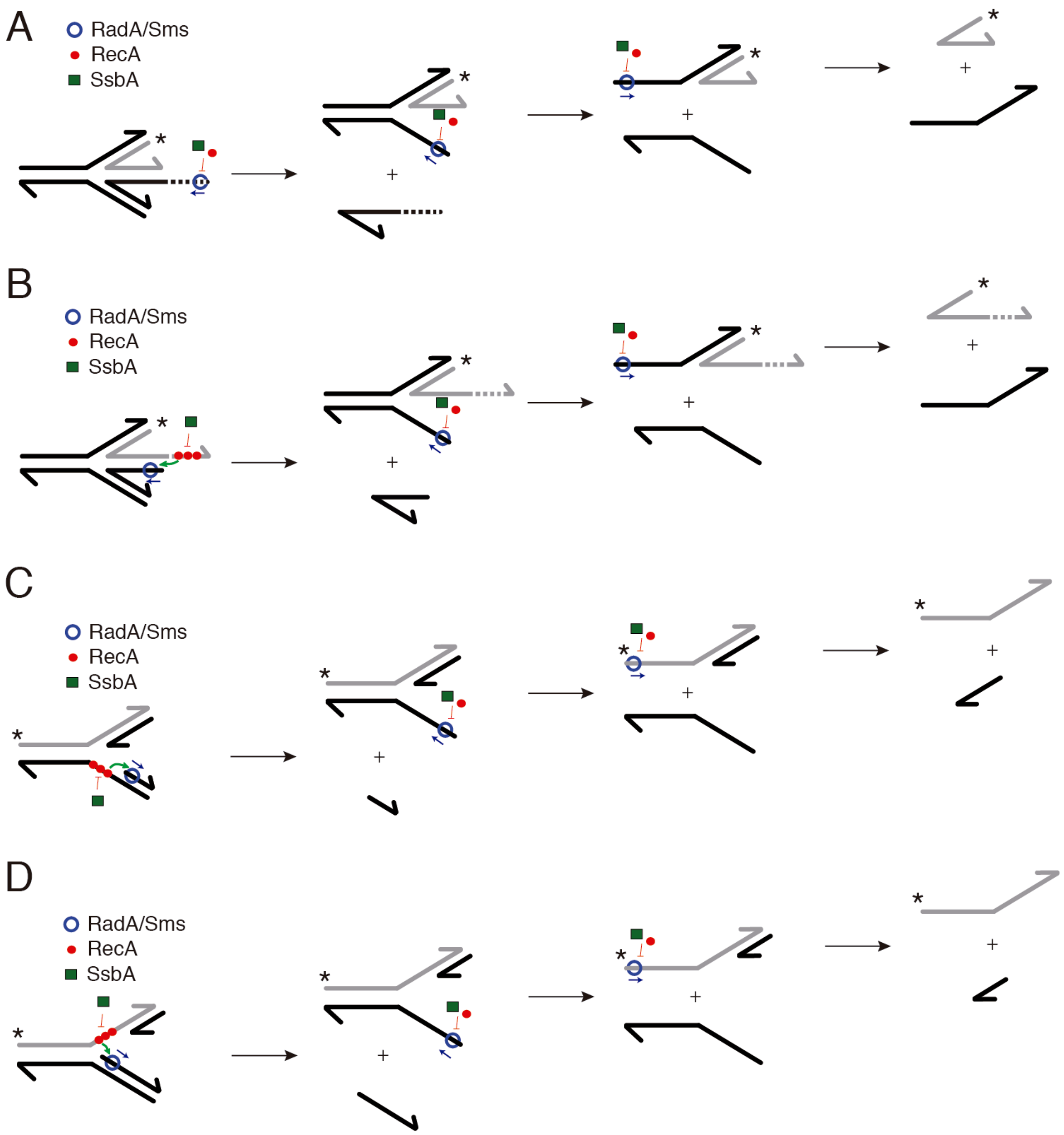


Figure S6. RecA action on RadA/Sms activity over reversed (A-B) and stalled forks (C-D). (A) RecA (red filled circle), or SsbA (green filled square) competes RadA/Sms (blue empty circle) for binding to the 5′ tails of the HJ-Lag DNA and its unwinding intermediates. RadA/Sms-mediated unwinding yields a 3-way junction, a 5′tailed duplex, and the radiolabelled strand. (B) RecA nucleates on the naked nascent leading-strand tail of the [$\gamma$-$^{32}$P]-HJ-Lead DNA, but SsbA inhibits RecA nucleation. RadA/Sms partially displaces SsbA and interacts with and loads RecA. Then RecA loads RadA/Sms on the nascent lagging-strand to unwind it. RecA, or SsbA competes RadA/Sms for binding to the 5′-tail of the 3-way junction. RadA/Sms unwinding yields a 5′tailed duplex, and the [$\gamma$-$^{32}$P]-nascent leading-strand product. (C-D) SsbA competes RecA for binding to the forked-Lag or forked-Lead DNA substrate. RadA/Sms partially displaces SsbA and interacts with and loads RecA onto the parental gap. Then, RecA loads RadA/Sms on the nascent lagging-strand to unwind it, to yield the cognate RadA/Sms substrate (3′-fork DNA), a 5′tailed duplex, and the radiolabelled strand.

Figure S7

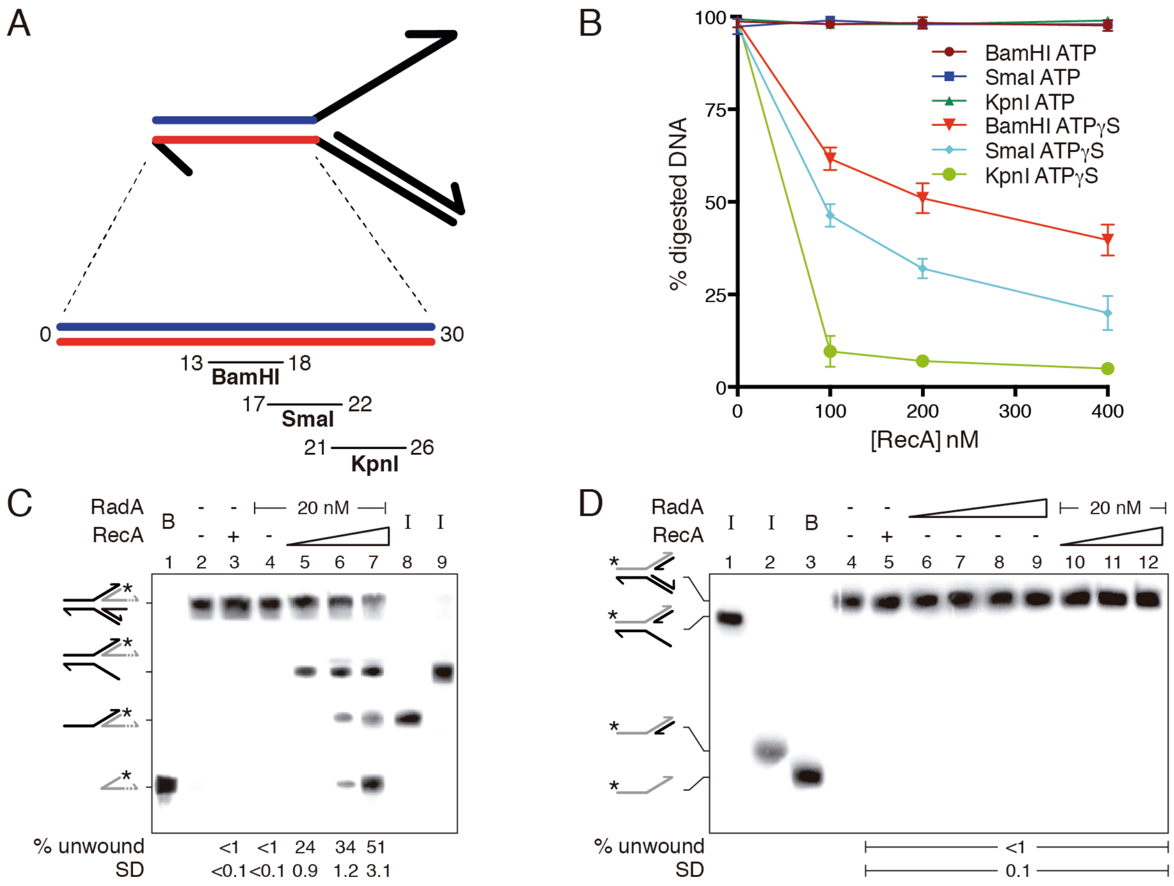


Figure S7. RecA does not remodel 5´-fork DNA and does not help RadA/Sms to unwind a fully replicated DNA (3´-5´-fork DNA). (A) The 5´-fork DNA substrate contains partially overlapping target sites for KpnI, SmaI and BamHI restriction enzymes at the depicted positions. (B) RecA (100-400 nM) was pre-incubated with the [$\gamma$-$^{32}$P]-5´-fork DNA in buffer A containing 400 $\mu$M ATP or ATP$\gamma$S for 15 min at 37 ºC. Then, the mixture was incubated with 1 unit of the KpnI, SmaI or BamHI restriction enzyme (60 min, 37 ºC). The samples were deproteinized and separated by 6% native PAGE. Gels were dried, visualized by phosphor imaging and the amount of digested DNA quantified. (C) The [$\gamma$-$^{32}$P]-HJ-Lead DNA substrate was preincubated with RecA·ATP$\gamma$S (100-400 nM) in buffer A (5 min, 30º C), then RadA/Sms (20 nM) and 2 mM ATP were added and unwinding was assayed (15 min, 30º C). The intermediates and products were separated after deproteinization of the samples. *, denotes the labelled strand; B, boiled product; half of an arrowhead, the 3´ends; -, no protein added; I, substrate intermediates. In lanes 8 and 9, the radiolabelled intermediates were loaded. The fraction of unwound products in three independent experiments was quantified, and the mean ± SD represented. (D) [$\gamma$-$^{32}$P] 3´-5´-fork DNA (0.5 nM in molecules) was incubated with the indicated protein(s) (fixed RecA [400 nM], or increasing RadA/Sms [10 - 80 nM], or fixed RadA/Sms [20 nM] and increasing RecA [100 - 400 nM]) in buffer A containing 2 mM ATP for 15 min at 30º C. Reactions were stopped, deproteinised and separated by 6% native PAGE. Gels were dried and visualised by phosphor imaging. *, denotes the labelled strand; half of an arrowhead, the 3´ends; -, no protein added; B, boiled product; I, substrate intermediates. The labelled intermediates and product were run in parallel as mobility markers (lanes 1-3). The fraction of unwound products in three independent experiments was quantified, and the mean ± SD represented.
